# Rho GTPases signaling mediates aggressiveness and differentiation in neuroblastoma tumors

**DOI:** 10.1101/2024.11.20.624451

**Authors:** María A. Gómez-Muñoz, Mónica Ojeda-Puertas, Luis Luna-Ramírez, Aida Amador-Álvarez, Ismael Rodríguez-Prieto, Juan Antonio Cordero Varela, Ricardo Pardal, Francisco M. Vega

**Author notes:** **Correspondence:** Dr Francisco M Vega, Department of Cell Biology, Faculty of Biology, Universidad de Sevilla. Avda. Reina Mercedes 6, Sevilla 41012, Spain. Dr Ricardo Pardal. Department of Medical Physiology and Biophysics, Universidad de Sevilla. IBiS. Avda. Manuel Siurot s/n, Sevilla 41013, Spain. NYU Grossman School of Medicine. Department of Pathology. Smilow Research Center, 10016, New York, USA.

## Abstract

**Background:** Neuroblastoma (NB) is a pediatric cancer with highly variable outcomes, necessitating improved understanding of the molecular pathways driving its progression. Intratumor cellular heterogeneity related to neural differentiation has emerged as a defining characteristic that can explain its aggressive behavior. Although recurrent driver mutations are not typically observed in these tumors, Rho GTPases signaling genes have been identified as one of the most frequently mutated in aggressive NB cases. Rho GTPases are key regulators of cell morphology, migration, and differentiation, yet their role in NB remains underexplored. This study aims to comprehensively evaluate the expression and clinical significance of Rho GTPase signaling networks in NB tumors.

**Methods:** We analyzed the expression profiles of Rho GTPases, their regulators, and effectors, across multiple NB patient cohorts. Gene expression correlations with clinical parameters were assessed, and bioinformatics analyses were employed to identify gene expression patterns and interactions in tumors. Functional studies were performed in NB cell lines and in vivo models to validate the role of key Rho GTPases, including Cdc42, in NB progression and differentiation.

**Results:** Our analysis revealed widespread dysregulation of Rho GTPase signaling in NB tumors. Specific GTPases, such as *RHOA* or *RHOV*, were upregulated in advanced disease stages, while others, including *RHOB*, *RHOU* and *CDC42*, were downregulated and associated with poor prognosis. A minimal Rho-related gene signature was identified as a strong predictor of NB patient survival. Functional validation highlighted Cdc42 as a key regulator of NB differentiation, where its downregulation was necessary for maintaining the malignant, undifferentiated phenotype of NB cells. We also identified *ARHGAP31*/CdGAP as a critical regulator of Cdc42 in NB progenitor cells, suggesting a mechanism for Cdc42 suppression in aggressive NB.

**Conclusions:** An important role for Rho GTPase signaling in NB progression is revealed, providing a foundation for further exploration of Rho GTPase-targeted therapies in NB. In particular, Cdc42 signaling intervene in the balance between differentiation and stemness in NB cells, suggesting specific signaling events controlling the identity and plasticity of NB cells.

## Background

The oncogenic transformation of sympathetic nervous system progenitors during embryonic neural crest development can lead to neuroblastoma (NB) tumors (1). NB is the most prevalent extracranial solid tumor during childhood, accounting for 15% of all pediatric oncology-related deaths. It is remarkably characterized by its variable clinical presentation, ranging from spontaneous regression to aggressive metastatic tumors resistant to multimodal therapy (2). According to recent single-cell expression studies and transcriptomic data (3–5), intratumoral cellular heterogeneity strongly contributes to NB presentation and outcome. Tumors in low-risk patients are characterized mostly by the presence of committed proliferative neuroblasts with a high degree of adrenergic differentiation; meanwhile, in high-risk tumors, an undifferentiated cell population with a mesenchymal phenotype and resemblance to neural crest stem cells can be identified (6). This latter cell population is thought to present increased resistance to chemotherapy and increased metastatic capacity (7–9).

As with most pediatric tumors, neuroblastoma initiation and progression do not typically involve recurrent mutations in common oncogenic drivers; instead, chromosomal rearrangements, including chromothripsis, are more frequent (10,11). Nevertheless, genomic and sequencing analyses have revealed that some discrete gene families are frequently found mutated in aggressive or relapsed neuroblastoma (10–12). This is the case for the extensive Rho family of small GTPases and their regulators. The Rho GTPases are a group of signaling molecules that function as molecular switches, alternating between an active GTP-bound form and an inactive GDP-bound form (13). This GTPase switch is positively modulated by guanine exchange factors (GEF) proteins and negatively regulated by GTPase activating factors (GAP) or guanine dissociation inhibitors (GDI). In addition, the activity of all Rho proteins is regulated by posttranslational modifications and gene expression (14). Furthermore, many members of this family, called atypical Rho GTPases (including RhoD, RhoF and Rnd1, 2 and 3), have virtually no GTPase activity and are regulated mainly at the transcriptional or posttranscriptional level. Once activated, Rho GTPases interact with different specific effectors that perform diverse cellular functions, which are mostly related to the control of cytoskeletal dynamics, cell movement, cell adhesion or cell proliferation. Rho proteins thus affect multiple cellular processes involved both in normal physiology and in disease, including development, inflammation and cancer progression (15). As essential modulators of actin and the microtubule cytoskeleton, Rho GTPases play important roles during neuronal development and differentiation (16). In particular, the Rho GTPases Rac1 and Cdc42 are involved in neuron development and axon growth (17,18), and Rho signaling has been described as pivotal in the regulation of neural crest cell fate and migration (19).

Here, we extensively describe Rho GTPase signaling expression patterns associated with different clinical parameters in NB patient tumors. We identified the dysregulation of some Rho GTPases in neuroblastoma tumors associated with malignancy and poor survival, providing new opportunities for treatment and prognosis. In particular, a correlation between low Cdc42 expression and malignancy in neuroblastoma tumors has been established, underscoring the crucial role of this Rho GTPase in neuroblastoma cell proliferation and differentiation. We propose that low levels of Cdc42, via the upregulation of its negative regulator *ARHGAP31*/CdGAP, could play a role in the self-maintenance of proliferative undifferentiated NB cells with malignant characteristics.

## Materials and methods

### Gene expression analysis in neuroblastoma patient tumors

Rho GTPase gene expression was analyzed via microarray and RNA-Seq data from neuroblastoma patient cohorts on the R2 platform ‘R2: Genomics Analysis and Visualization Platform (http://r2.amc.nl (accessed on June 15, 2022))’. Databases and included Rho GTPase regulators and effectors are summarized in Additional Tables S1 and S2. Gene selection for Rho GTPases regulators and effectors was done on the basis of literature (at least some published results with connection with some Rho GTPase) and relevant expression in the used NB patient cohorts. The data were managed and processed with Excel, Prism and RStudio (*ggplot2*, *circlize* y *ComplexHeatmap* packages) and displayed via box plots, violin plots or heatmaps. Kaplan‒Meier survival analyses were performed via the R2 platform-by-scan method with Bonferroni-corrected p values.

Gene Ontology, pathway enrichment, and protein interaction analyses were conducted via the Reactome (https://reactome.org/), Gene Ontology (http://geneontology.org), and STRING (https://string-db.org) platforms. Data representation was performed with RStudio (*ggplot2* package). For STRING analysis, a minimum interaction score of 0.04 and a maximum of 10 interactors were used. Unsupervised stratification of patient tumor samples by Rho GTPase expression was performed using K-means clustering on the R2 platform, with 3 groups and 10 rounds.

### Correlated expression analysis and WGCNA

Expression correlation studies between the Rho GTPase family of genes and their regulators were performed by calculating the Pearson correlation coefficient, with data from GSE62564, and represented by a heatmap in RStudio with the packages ComplexHeatmap, ggplot2 and circlize. Weighted correlation network analysis (WGCNA) was performed using the GSE62564 dataset in RStudio. Jaccard analysis was performed with the RStudio *ggplot2*, *circlize* and *ComplexHeatmap* packages.

### Cox regression analyses for survival

A gene expression signature was built from the training dataset (TARGET) according to the expression of Rho GTPases and their regulators, via multivariate Cox regression. A Lasso penalty was applied to minimize the risk of overfitting using the *glmnet* package. The penalty parameter was calculated using a tenfold cross-validation (at the minimum partial likelihood deviance) of the complete cohort with one thousand iterations, randomly dividing 90% and 10% of patients into training and testing sets, respectively. Risk scores were calculated for each patient by multiplying the expression of each gene by its corresponding Cox regression prognostic index.

Cox proportional hazards regression analysis was performed to obtain a gene expression signature using a training dataset (TARGET) according to the expression of Rho GTPases and their regulators.

Survival was predicted at multiple times using a Cox proportional hazards regression model from the obtained risk scores to draw receive operator curves (ROC) and compute the area under the curve (AUC). Considering the median value of the risk score as a threshold, patients were classified into two groups, low -risk and high-risk, and a Kaplan-Meier survival analysis was performed. Univariate Cox regression, including the expression signature and several clinical parameters, was performed to preselect a set of significant variables. An additional validation step in another patient cohort (GSE62564) was performed by following the same procedure, using the already calculated coefficients for the signature.

### Cell culture and transfection

The human NB cell line SK-N-SH was obtained from the American Type Culture Collection (ATCC) and cultured in DMEM + GlutaMAX (GIBCO BRL) supplemented with 10% fetal bovine serum (FBS) (GIBCO BRL), 100 µg/ml streptomycin, and 100 U/ml penicillin. CHLA 20 and CHLA 255 cell lines were obtained from the COG Cell Line and Xenograft Repository (Texas Tech University Health Sciences Center, USA) and cultured in IMDM+GlutaMAX with 15% FBS, 100 µg/ml streptomycin, 100 U/ml penicillin and 1X ITS (5 µg/ml insulin, 5 µg/ml transferrin and 5 ng/ml selenium acid). All cells were maintained at 37°C in a 5% CO2 humidified atmosphere. Patient-derived xenograft (PDX) tumors were mechanically and enzymatically dispersed to generate NB48T cell cultures (9).

For transfection, cells at 80-90% confluence were transfected with 2.5-5 µg of plasmid DNA (pcDNA3-EGFP-Cdc42-wt, Addgene #12975) or 50-100 nM siRNA (CdGAP Smart pool or Cdc42 siRNA, Horizon) using Lipofectamine 2000 (Life Technologies). EGFP-Cdc42 expressing cells were selected with Geneticin (G418) for 7 days and sorted by GFP expression.

For undifferentiated tumorsphere formation, SK-N-SH or NB48T cells were cultured in neural crest media (DMEMF-12 medium supplemented with 1% N2, 0.5% B27, 15% FBS, 100 µg/ml streptomycin, 100 U/ml penicillin, 10 μg/ml FGF, 20 μg/ml IGF-1, and 20 μg/ml human EGF) at low density in low-adherence plates as previously described (20). Cell viability was measured via AlamarBlue (Thermo Scientific).

For differentiation assays, NB cells were seeded in complete medium and treated 24h after with standard culture media (DMEM: F12 + 100 μg/ml streptomycin + 100 U/ml penicillin + 1% N2 supplement + 2% B27 supplement with vitamin A) or media containing 10 µM all-trans retinoic acid (ATRA) for 7-10 days, as described in (9). Cell Profiler software (21) was used for cytoskeleton-based morphological profiling and differentiation analysis. Briefly, cells seeded at low density were fixed, stained with fluorescent phalloidin and DAPI, and imaged with an Olympus BX-61 microscope. An image analysis pipeline corrected illumination, segmented cells, and extracted morphological features. Relative z scores of morphological variables such as form factor or number of cell branches, were represented for comparison across cell populations.

### Fluorescence-activated cell sorting (FACS)

Cell sorting was done using a BD FACS Jazz sorter as previously described (20). NB cells were detached with versene, resuspended in FACS medium (L15 + 2 mg/ml BSA, 1% HEPES, 1% penicillin/streptomycin), and incubated with CD44 primary antibody (BD Pharmingen, mouse IgG2b) for 40 min at 4°C. Cells were then incubated with an Alexa 488-conjugated secondary antibody (1:1000, Life Technologies) for 30 min on ice, strained through a 40 µm filter, and sorted into CD44hi or CD44-populations.

### Immunofluorescence and immunohistochemistry

For immunofluorescence, cells were fixed with 4% PFA in PBS for 15 minutes, permeabilized with 0.2% Triton X-100 in PBS for 10 minutes at 4°C and blocked with PBS + 1% BSA for at least 1 hour. Primary antibodies used were: CD44 mouse (1:200 BD Pharmingen), Cdc42 rabbit (1:1000 Cell Signaling), Ki67 rabbit (1:200 Thermo Scientific), Nestin rabbit (1:1000 Millipore), Nestin mouse (1:1000 R&D Systems), α-tubulin mouse (1:1000 Sigma), acetylated tubulin mouse (1:1000, Santa Cruz), Tuj1-βIII tubulin mouse (1:1000 abcam) and CdGAP (1:500, Santa Cruz). The corresponding fluorescently labeled secondary antibodies (Alexa Fluor 488, 568 or 633) (Life Technologies) were used at a 1:1000 dilution. Actin was stained with Alexa Fluor 488 or 568 phalloidin (1:500) (Life Technologies) and nuclei with 4’,6-diamino-2-phenylindole (DAPI) 1 μg/ml.

For immunohistochemistry, paraffin-embedded tissue sections were hydrated, subjected to antigen retrieval with citrate buffer (pH 6), and incubated with the following primary antibodies: DDC rabbit (1:1000, Millipore), Ki67 rabbit (1:200, Thermo Scientific), and VRK1 rabbit (1:500, Sigma). Secondary antibodies used were biotinylated anti-rabbit and anti-mouse (1:2000, Vector Lab.). The Vectastain ABC kit and DAB peroxidase substrate (Vector Laboratories) were used for staining. Images were analyzed with QuPath software (22).

### Gene expression by RT-PCR

RNA was extracted using the QIAamp RNeasy Mini Kit (Qiagen). Preamplification, when needed, was performed with the Affymetrix Whole Transcript Expression Array Kit (Ambion). For reverse transcription, 1 μg of RNA was converted to cDNA using qScript (QuantaBio). For RT-qPCR, Fast Sybr Green polymerase (Applied Biosystems) was used with glyceraldehyde-3-phosphate dehydrogenase (GAPDH) used as the control. Primers (0.2 μM, see Additional Table S3) and 50-300 μg of genetic material were used in a 7500 Fast Real Time PCR System (Applied Biosystems) in 96-well plates. Data analysis was performed using the ΔCt (cycle threshold) comparative method.

### Western blot and GTPase activity assays

Cells were lysed with buffer (50 mM Tris-HCl (pH 8), 0.5 mM EDTA, 150 mM NaCl, 1% Triton X-100, protease, and phosphatase inhibitor cocktails). Western blotting was performed using primary antibodies overnight and HRP-conjugated secondary antibodies for 1 hour. The following antibodies were used: Cdc42 (1:500, Cell Signaling), GFP (1:500, Invitrogen), α-tubulin (1:2000, Sigma), GAPDH (1:2000, Trevigen), CdGAP (1:500, Santa Cruz), and HRP secondary antibodies (1:1000, Jackson ImmunoResearch). Detection was performed with ECL reagent (Thermo Scientific) and a ChemiDoc™ Touch Imaging System (BioRad). Protein quantification used FIJI software (National Institutes of Health).

For the GTPase activity assay, cells were lysed in pull-down buffer (50 mM Tris-HCl, pH 7.2, 0.5 M NaCl, 5 mM MgCl2, 1% Triton X-100, 0.1% SDS, protease, and phosphatase inhibitor cocktails). Lysates (800-1000 μg) were incubated with GST-PAK Sepharose beads (8-10 μg) for 1 hour at 4°C. The controls included incubation with 100 μM GTPγS or 10 mM GDP (Sigma). Bound proteins were resolved by electrophoresis and detected with the specified antibodies.

### Quantitative label-free proteomics (LFQ)

For LFQ proteomics, samples were processed using the FASP DIGESTION protocol with Millipore Microcon 30K filters (Merck). Proteins were reduced with 15 mM tris(2-carboxyethyl)phosphine (TCEP) and alkylated with 30 mM chloroacetamide (CAA) in urea, incubated at 37°C for one hour in the dark. Digestion was done with trypsin (Promega) at a 1:20 enzyme-to-protein ratio at 37°C for 12 hours. Peptides were analyzed using a nano HPLC system (nano LC 1000) coupled to a Q Exactive Plus Orbitrap mass spectrometer (Thermo Scientific) with a nanospray ionization source (Proxeon Biosystems). The data were processed with Proteome Discoverer software (Thermo Scientific) using an FDR<1% and MetaboAnalyst software (23). Protein interaction and functionality were modeled using the STRING platform.

### Experimental tumor assay

Subcutaneous tumors were induced by injecting 1×10^5^ to 1×10^6^ cells in 100µl PBS:Matrigel (1:1) (Corning) into C.B-17 SCID (Severe Combined Immunodeficiency.:C.B-17/IcrHan HsdPrkdcscid) mice. Tumor development was monitored biweekly until tumor onset, then weekly for 2 months or until the tumors reached 10mm across.

For orthotopic tumor formation, 2×10^4^ luciferase-expressing NB cells were injected with a Hamilton needle into the adrenal gland of C.B-17 SCID mice. Mice were anesthetized and a 1 cm incision in the retro-lateral abdomen provided access for injection. After injection, mice were monitored for 3 days for recovery and then imaged weekly using an IVIS Spectrum system (Xenogen) after anesthesia with isoflurane and luciferin retroperitoneal injection (150 mg/kg). Mice were observed for up to 2 months and sacrificed upon excessive tumor growth or lower lumbar swelling.

### Statistical analysis

Statistical parameters, including the exact n value, and the mean ± S.D. or ± S.E.M. are described in the figures and figure legends. All statistical analysis were performed using the GraphPad Prism 9.5.1 software. Data represented on boxplots are representing the 10th-90th percentile distributions and medians. For statistical comparisons between two experimental groups a nonparametric t-test was performed. For the analysis and comparison of data represented as growth curves, linear regression and curve comparison were performed. Normalization by z score was used for the analysis of intensity data and cellular profiling. Experiments were independently repeated as described in the figure legends.

## Results

### Rho GTPase signaling is dysregulated in NB tumors

Given the relevance of Rho GTPase signaling in cancer progression, neural development and differentiation, and its identification as one of the pathways associated with the greatest number of mutations in NB tumors, we analyze the expression of these pathways in detail in human NB patient tumor cohorts. We followed an analysis pipeline designed to identify Rho-related signaling clusters correlated with NB aggressiveness, and identified candidate genes for functional validation (Fig. 1a)

**Fig. 1.**
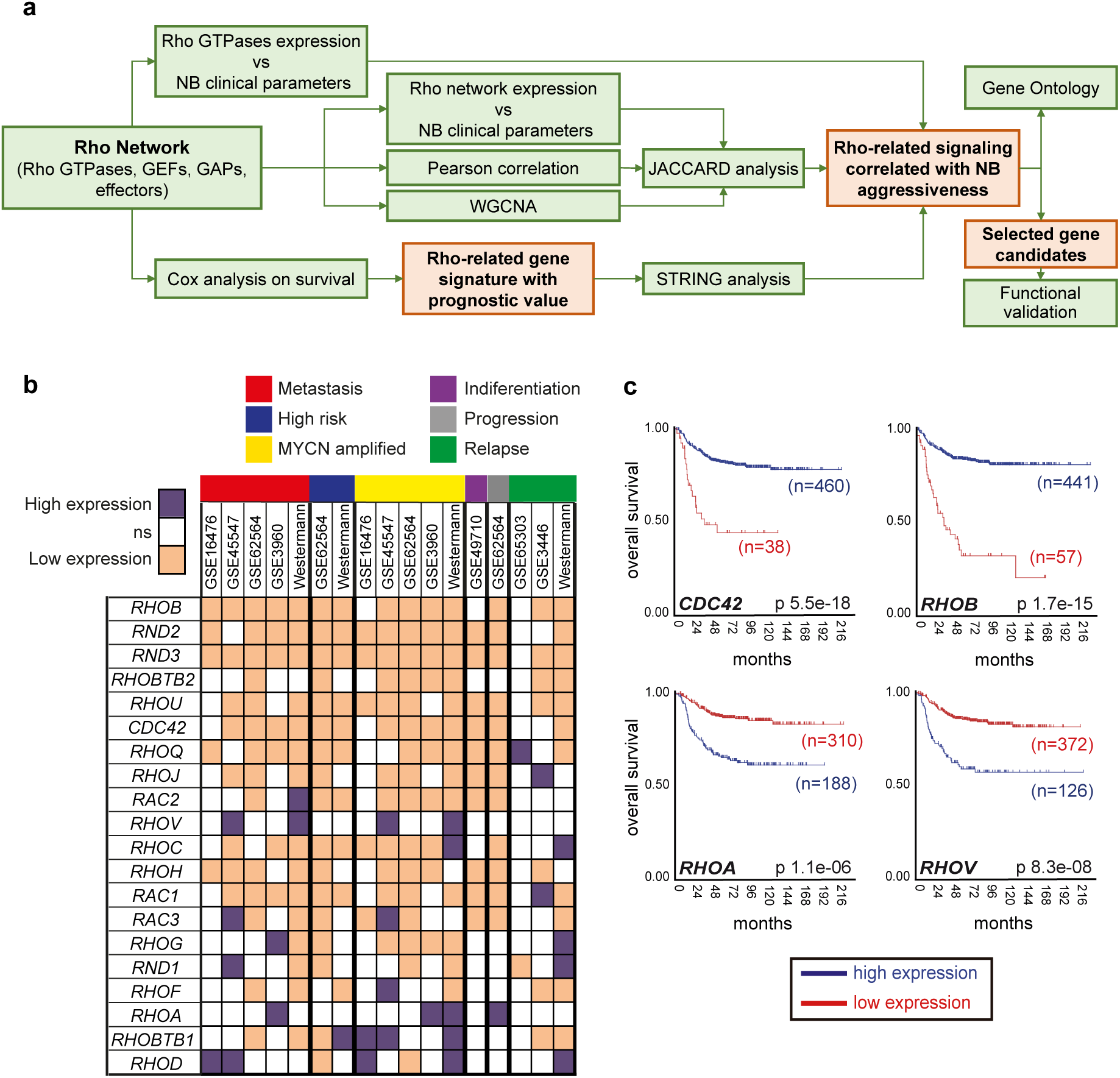
The expression of the Rho GTPase family is dysregulated in NB tumors and predicts outcome. **a** Schematic illustration of the in-silico analyses workflow followed for the Rho GTPase network expression studies in NB patient’s datasets. **b** Table summarizing Rho GTPases expression significance in relation to different NB clinical parameters. The parameters are indicated with a different color code. High and low expression indicates a significant correlation of a high expression or low expression respectively of the Rho GTPase with the clinical variable. NS: not significant. **c** Kaplan-Meier overall survival analysis of representative Rho GTPases. Data from cohort GSE62564. n= number of samples. Bonferroni adjusted p values are shown.

We first analyzed the expression of all 20 Rho GTPases and their correlations with clinicopathological features, such as INSS stage, MYCN status, risk, histology, metastasis or relapse. Using different expression databases from patient sample cohorts, we detected significant deregulation of the expression of some GTPases during tumor progression (Fig. 1b, Additional Figs. S1 and S2). There is a group of GTPases, including *RHOB*, *RND2*, *RND3*, *RHOBTB2*, *RHOU*, *CDC42*, *RAC2* and *RHOH*, whose expression inversely correlates with malignant clinical parameters. Conversely, increased expression of the Rho GTPases *RHOA*, *RHOV* or *RHOD* is correlated with several features of tumor progression. The results for the rest of the GTPases were variable, depending on the patient cohort or parameter analyzed. K-means stratification of patient tumor samples according to Rho GTPase expression revealed a subset of stage 4, *MYCN* amplified tumors with consistent downregulation of the *CDC42*, *RND2* and *RND3* genes and upregulation of the *RHOA*, *RHOG* and *RHOBTB1* genes across datasets (Additional Fig. S3).

Moreover, Kaplan-Meier survival analysis determined that the expression of some Rho GTPases was sufficient to stratify NB patients according to their overall survival. High expression of the GTPases *RHOA*, *RHOV* and *RHOBTB1* or low expression of *RHOB*, *RHOC*, *RHOD*, *RHOH*, *CDC42*, *RHOJ*, *RHOQ*, *RAC2*, *RND2* and *RHOU* strongly associated with poor patient overall survival (Fig. 1c and Additional Fig. S4).

Given that the majority of alterations in Rho GTPase signaling in cancer comes from changes in their regulators, we analyzed the expression of an ample set of genes from the Rho signaling network in NB patient samples and their correlation with clinical parameters. On the basis of the known literature and relevant expression in NB patient cohorts, we selected a group of GAPs (n=51), GEFs (n=55), GDIs (n=4) and effectors (n=26) (Additional Table S2). We first explored the individual expression of every gene in patient cohorts and their coexpression with the Rho GTPases, generating hierarchical heatmaps of the Pearson correlation and identifying clusters of genes whose expression changed together (Fig 2a and Additional Table S4). We subsequently performed a weighted correlation network analysis (WGCNA) to define groups of genes whose expression changes correlated with particular clinical parameters, such as tumor stage, MYCN status or age at diagnosis (Fig. 2b and Additional Table S4). Finally, we followed a Jaccard similarity analysis to combine both approaches and better define gene circuits including Rho GTPases and potential regulators whose coexpression associates with relevant clinical parameters. Following this approach, we selected 3 relevant groups of genes whose expression correlates in tumors, and associates similarly with clinical parameters (Fig. 2c and Additional Table S4). Group 1 includes the GTPases *RHOA*, *RHOC*, *RAC2*, *RHOG* and *RHOH*. Group 2 includes among others the GTPases *CDC42* and *RAC1*, together with different specific regulators and effectors, such as *ARHGEF7*, *DOCK3*, *DOCK4* and *PARD3B*, and is negatively associated with adverse clinical parameters. Group 3 includes relevant effectors such as *TIAM1* and *ROCK1* and is associated with malignancy, although the number of genes in this group was very low. Gene ontology analysis performed with these genes relevantly showed an association of Group 2 genes with neural development, proliferation and differentiation (Fig. 2d and Additional Table S4). Moreover, Reactome pathway analysis identifies p75 NTR receptor-mediated signaling, semaphorins, axon guidance and nervous system development, and Ephrin signaling among the most relevant pathways overrepresented (Additional Table S4). The analysis of genes in group 1 revealed that GPCR signaling was a specific signaling pathway for this group.

**Fig. 2.**
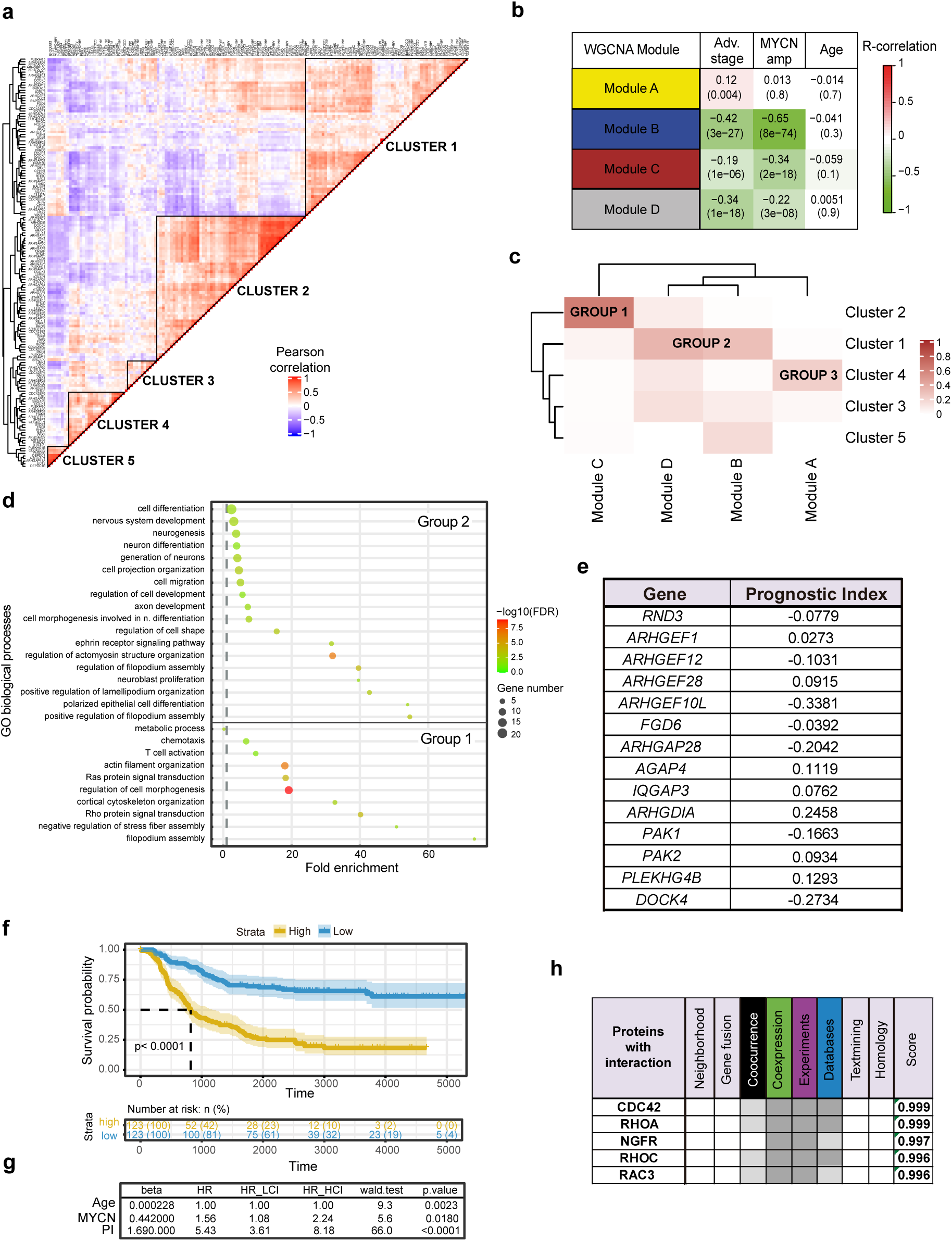
Rho GTPase network genes expression correlate with NB clinical behavior. **a** Correlation between genes from the Rho GTPase network in NB tumor patient data. Heatmap colored according to Pearson correlation. Clusters with genes with high correlation in tumor samples are shown. **b** Modules resulting from the Weighted Correlation Network Analysis (WGCNA) containing genes from the Rho GTPase network. Genes in the same module are coexpressed and hold a significant correlation with the indicated NB clinical parameters (MYCN amplification, INSS stage, or Age at diagnosis). Upper value indicates the correlation value R. Lower value indicates p value. **c** Jaccard analysis performed with the gene clusters from A together with the gene modules from B. Distances between groups were estimated and represented in a heatmap. The selected 3 groups of relevant genes resulted from the combination of both analyses. **d** Gene ontology enrichment analysis results showing the most relevant biological processes associated with the genes in Group 1 and Group 2 obtained from c. Data is from GSE45547 dataset. The complete list of genes in each cluster, module or group can be seen on Additional Table S4. **e** Rho-related gene signature with predictive capacity for NB patient survival. The associated prognostic index (PI) obtained from Cox regression analysis is shown. **f** Kaplan-Meier analysis using Rho family prognostic index (PI). Samples with higher PI values have lower survival than the group of samples with lower PI (Strata: classification of patients in survival data according to PI value, PI High = high PI values, higher risk disease, PI Low = low PI values, lower risk disease). **g** Hazard ratio (HR) obtained by performing a Univariate Cox Regression Analysis for the PI index and other clinical parameters in relation to overall survival (p value = likelihood ratio test, Wald test and logRank test). Data from cohort TARGET. **h** Overall likelihood score of interaction with the proteins encoded by the gene signature according to STRING analysis, stratified by different factors.

Taken together, these analyses show a relevant and complex deregulation of the Rho GTPase signaling network in NB tumors associated with clinical progression and differentiation status.

### A Rho-related gene expression signature predicts survival in neuroblastoma patients

Given the deregulation of the Rho signaling network observed in neuroblastoma tumors and its association with neuroblastoma patient survival and malignancy, we performed Cox proportional hazards regression analysis to identify a specific minimal gene expression signature predictor of survival. We initially used the TARGET patient cohort dataset. These analyses described the expression of the genes *RND3*, *ARHGEF1*, *ARHGEF12*, *ARHGEF28*, *ARHGEF10L*, *FGD6*, *ARHGAP28*, *AGAP4*, *IQGAP3*, *ARHGDIA*, *PAK1*, *PAK2*, *PLEKHG4B* and *DOCK4* as the minimal Rho-related expression signature with prognostic value in neuroblastoma (Fig. 2e). Every gene has an associated prognosis index (PI), which corresponds to their contribution to overall survival prediction. Area under curve (AUC) analysis shows values over 0.74 for the prediction at different time points (Additional Fig. S5a). The expression of the signature significantly stratified patients with worse survival (p<0.0001) (Fig. 2f). Moreover, when the ability of the Rho-related signature to predict outcome was compared with that of other clinical variables, the results show a similar or better performance of the former (Fig. 2g). The signature was validated in silico using the GSE62564 cohort, with similar performance values (Additional Fig. S5b-d).

The genes included in the signature were all regulators of Rho GTPases with the exception of *RND3*. We performed a STRING analysis to identify the most likely interaction relationships of these genes with Rho GTPases, showing Cdc42 and RhoA as the GTPases most likely affected (Fig. 2h and Additional Fig. S5e).

### Cdc42 expression counteracts proliferation and is associated with differentiation in NB cell lines

The Rho GTPase Cdc42 has been implicated in neuronal development and differentiation. We have shown that a low expression of *CDC42* is specifically associated with malignancy in neuroblastoma, including relapse, metastasis and low survival. Furthermore, the results of our Rho-related in silico expression analysis and the predictor gene signature, point to the Cdc42 GTPase pathway as an important mediator in neuroblastoma progression. To functionally validate our results, we decided to explore in more detail the roles of Cdc42 in neuroblastoma progression and malignancy. Thus, we overexpressed a tagged version of wild type Cdc42 in NB cell lines with low Cdc42 expression and analyzed the phenotypic changes in these NB tumor cells (Fig. 3).

**Fig. 3.**
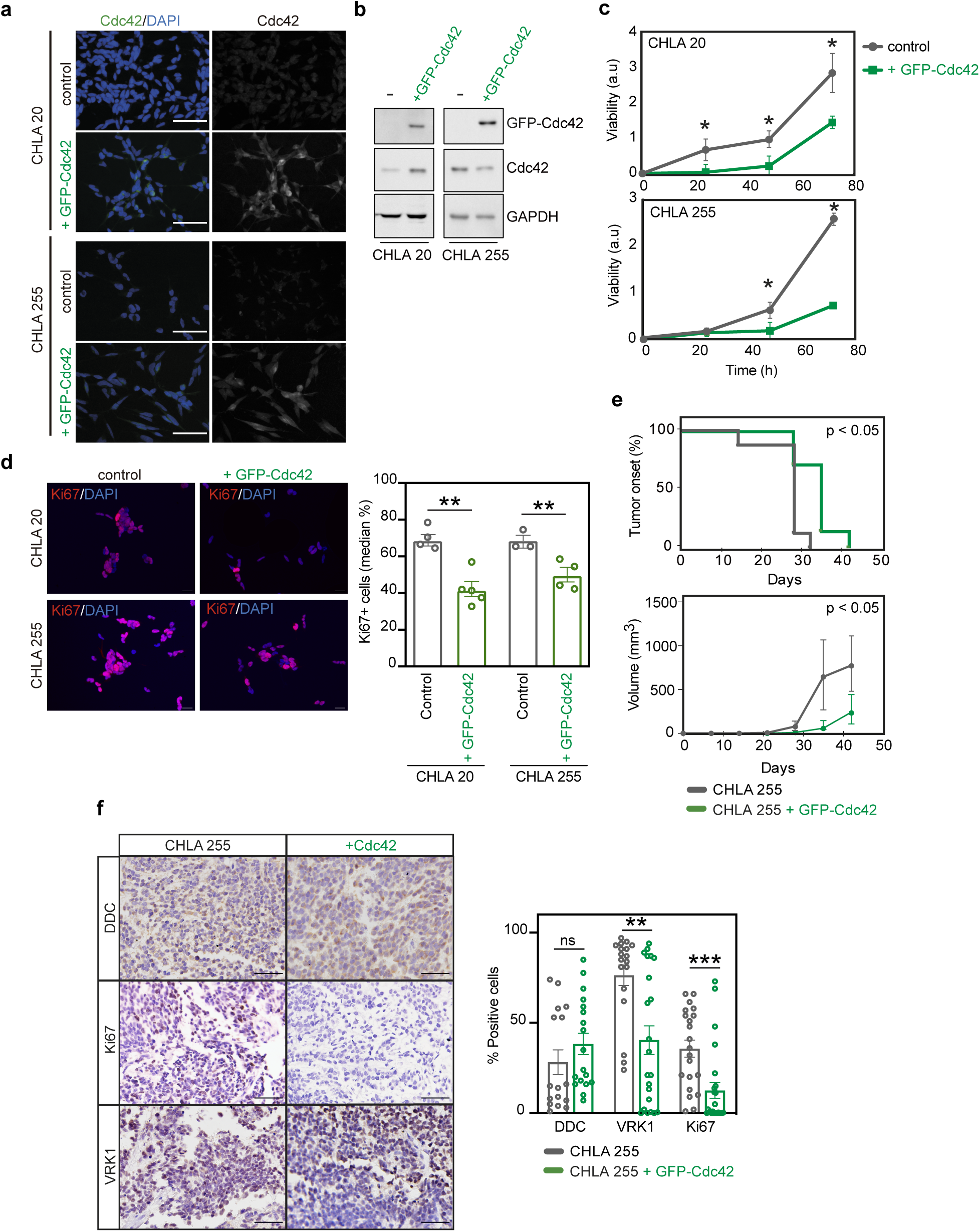
Cdc42 re-expression decreases proliferation in NB cells *in vitro* and *in vivo*. **a** Immunofluorescence showing Cdc42 staining in CHLA 20 and CHLA 255 cell lines with or without the expression of GFP-Cdc42. Scale bar = 50 μm. **b** Western blot showing Cdc42 and GFP-Cdc42 expression in the same cell lines. **c** Cell viability in CHLA 20 and CHLA 255 cell lines, with or without GFP-Cdc42 expression, over 72 hours (***=p<0.001, ****=p<0.0001, n=4). **d** Quantification of proliferating Ki67 expressing cells in CHLA20 cell line with or without the expression of GFP-Cdc42. Graph show mean +/−SD. Scale bar = 20μm. (**=p<0.01, n=4). **e** Analysist of tumor onset and growth in CHLA 255 xenografts with or without the expression of GFP-Cdc42. (n=10 per condition). LogRank statistical analysist significance is shown. Mean +/− SD is plotted in cell growth graph. **f** Immunohistochemistry from tumors described in E showing the expression of the indicated markers. Quantification shows the % of positive cells in each condition (mean +/− SD, **=p<0.01, ns= not significant). Scale bar = 50 μm.

Cdc42 re-expression was accompanied by an increase in Cdc42 activity in the cells (Fig. 3a and b and Additional Fig. S6a and b) and provoked a decrease in cell proliferation in vitro (Fig. 3c and d). When injected in immunocompromised mice, both in the flank and orthotopically, GFP-Cdc42 expressing cells produced less proliferative and smaller tumors, which appeared later (Fig. 3e and f and Additional Fig. S6c and d).

NB cells with overexpression of Cdc42 presented a morphology with increased neuronal-like features, even more accentuated after the induction of differentiation (Fig. 4a). Moreover, Cdc42 expressing cells tend to have more neurite-like protrusions enriched in α-tubulin aggregates (Fig. 4b). Cdc42 has been implicated in neurite outgrowth and microtubule polymerization (18). Microtubules enriched in the acetylated form of tubulin have been associated with stable neurite processes and neuronal differentiation, and they are only found in the perinuclear area in the case of undifferentiated cells (24,25). GFP-Cdc42 expressing cells presented an accumulation of the acetylated form of tubulin in neurite-like protrusions, indicating a shift towards differentiated cells (Fig. 4c and d).

**Fig. 4.**
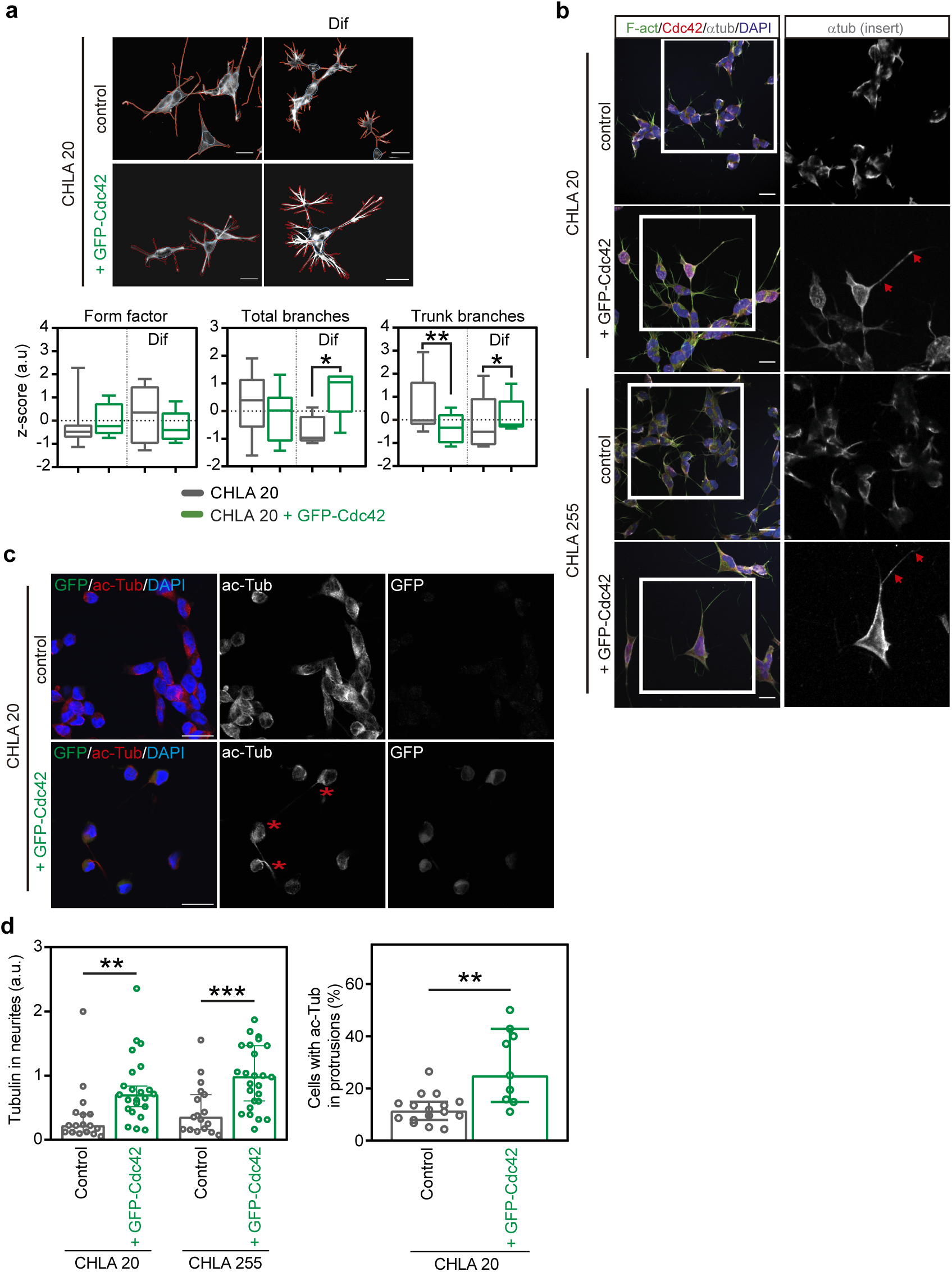
Cdc42 re-expression in NB cells drives a cytoskeletal reorganization associated with neuronal differentiation**. a** Morphological characterization of CHLA 20 cells overexpressing GFP-Cdc42 under standard culture conditions or differentiated conditions (Dif). Cell segmentation on the cell contour is shown in red. Quantification of morphological features, indicative of differentiation, are shown. *: p<0.05; **: p<0.01. Scale bar = 20 μm. **b** Cytoskeleton immunofluorescence in CHLA 20 and CHLA 255 cells with or without GFP-Cdc42 expression. Examples of tubulin accumulation in cell protrusions are indicated with a red asterisk. Scale bar = 20 μm. **c** Immunofluorescence for GFP, acetylated tubulin, and DAPI in CHLA 20 cells with or without GFP-Cdc42 expression. The presence of acetylated tubulin in cell extensions is indicated with red asterisks. Scale bar=20 μm. **d** (Left panel) Quantification of the relative abundance of tubulin foci, relative to the total number of cells from images like the ones shown in B. (a.u = arbitrary units, **=p<0.01, ***=p<0.001). (Right panel) Quantification of the cells positive for acetylated tubulin staining in cellular protrusions (Mean ± SEM, **=p<0.01).

To further demonstrate the association of Cdc42 with NB differentiation, we measured Cdc42 expression under different conditions showing that the expression of Cdc42 was increased in NB cells after differentiation treatment with all-trans retinoic acid (ATRA), both in SK-N-SH NB cells and PDX derived cells (Fig. 5a). We have previously described that the expression of high levels of the adhesion protein CD44 on tumorsphere-growing conditions selects for an undifferentiated malignant subpopulation of NB cells (9). Cdc42 expression was decreased in CD44hi cells and undifferentiated tumorspheres, compared to the CD44-counterparts (Fig. 5b). Moreover, Cdc42 expression inversely correlates with the neural crest undifferentiated marker Sox9 in NB tumor samples, and a low expression of Cdc42 was essential to detect patients bearing Sox9 positive undifferentiated tumors with worse outcome, revealing the particular downregulation of Cdc42 in undifferentiated stem-like cell populations (Fig. 5c). We have also grown tumorspheres from SK-N-SH cells, either overexpressing Cdc42 (Fig. 5d), or treated with an siRNA against Cdc42 (Fig. 5e). The results show an enrichment in differentiation marker expression (Tuj1 and TH) in the case of Cdc42 overexpression, and an increase in undifferentiated CD44hi cells in the case of Cdc42 silencing. Lastly, a label free proteomic analysis showed downregulation of Nestin and Vimentin, filament proteins associated with undifferentiation in the neural crest lineage, in CHLA20 cells overexpressing Cdc42, and upregulation of the neuronal marker NCAM (Fig. 5f). These results situate Cdc42 as an important modulator of NB differentiation whose downregulation is necessary for the maintenance of undifferentiated neural crest stem-like cell populations.

**Fig. 5.**
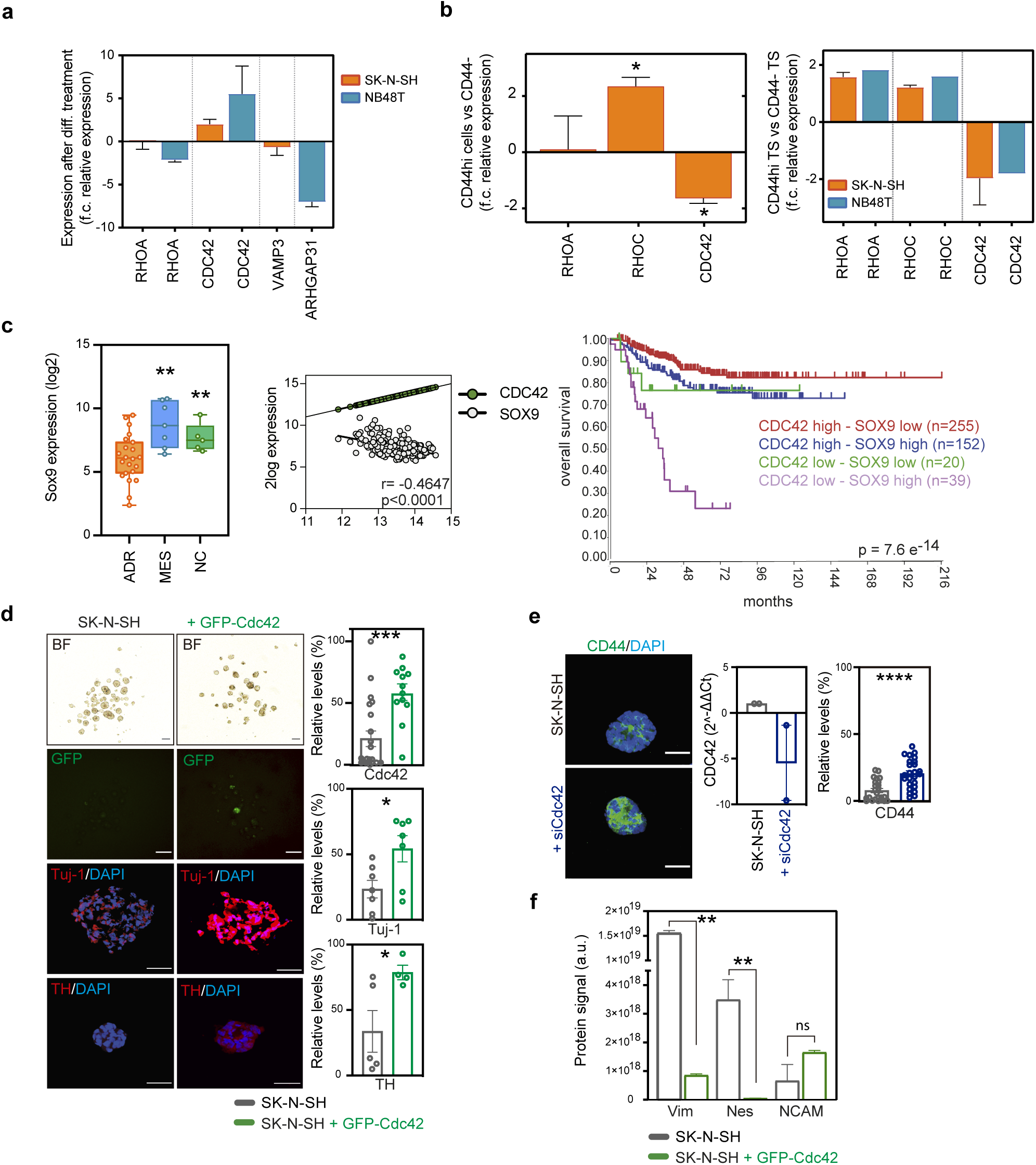
Cdc42 downregulation associates with an undifferentiated NB cell population. **A** mRNA expression of the Rho GTPases *RHOA* and *CDC42*, and the Cdc42 GAPs *VAMP3* and *ARHGAP31*, in the SK-N-SH and PDX NB48T cell lines before and after differentiation with ATRA (f.c = fold change, n = 3). **b** Quantification of the expression of the Rho GTPases *RHOA*, *RHOC* and *CDC42* in CD44hi sorted NB cells (left panel) or secondary tumorspheres derived from CD44hi NB cells (right panel), compared with their CD44 negative counterparts (f.c = fold change, n = 3). **c** Expression of *SOX9* in adrenergic tumors, mesenchymal tumors or neural crest tissues (left panel). Linear regression analysis of the expression of *SOX9* and *CDC42* in NB patient tumors (middle panel). Kaplan-Meier survival curves stratified by *CDC42* and *SOX9* gene expression combined (right panel). Long rank test p value is shown. Datasets: GSE90803 and GSE45547. **d** representative images from tumorspheres generated from SK-N-SH cells with or without GFP-Cdc42 expression. Relative expression levels of the indicated markers are quantified. Graphs show mean ± SEM. (*=p<0.05, ***=p<0.001). Scale bar= 200 μm (BF) or 50 μm (IF). **e** *CDC42* and CD44 expression in tumorspheres generated from SK-N-SH cells treated with an siRNA against Cdc42 or control. Scale bar = 50μm. (****=p<0.0001). **f** Results from label free proteomics analysis showing detection of the proteins Vimentin, Nestin and NCAM in SK-N-SH cells with or without GFP-Cdc42 expression. (**=p<0.01)

### The GAP *ARHGAP31*/CdGAP regulates NB cell neuronal differentiation by altering Cdc42 levels

We have stablished that the level of Cdc42 dictates cellular differentiation in NB tumor cells and that malignant NB stem-like cells decrease Cdc42 to maintain their undifferentiated status. Searching for Cdc42 regulators whose expression could be specifically altered in undifferentiated NB cells, we identified that *ARHGAP31*/CdGAP (Cdc42 GAP) is overexpressed in NB patient tumors with mesenchymal character or neural crest tissues (Fig. 6a and Additional Fig. S7a), and is downregulated in differentiated NB cells and upregulated in undifferentiated tumorspheres (Fig. 5a and Fig. 6b and c). CdGAP has been described as a negative regulator of Cdc42 and Rac1 activities, and changes in its expression has been proposed to mediate Cdc42 function (26).

**Fig. 6.**
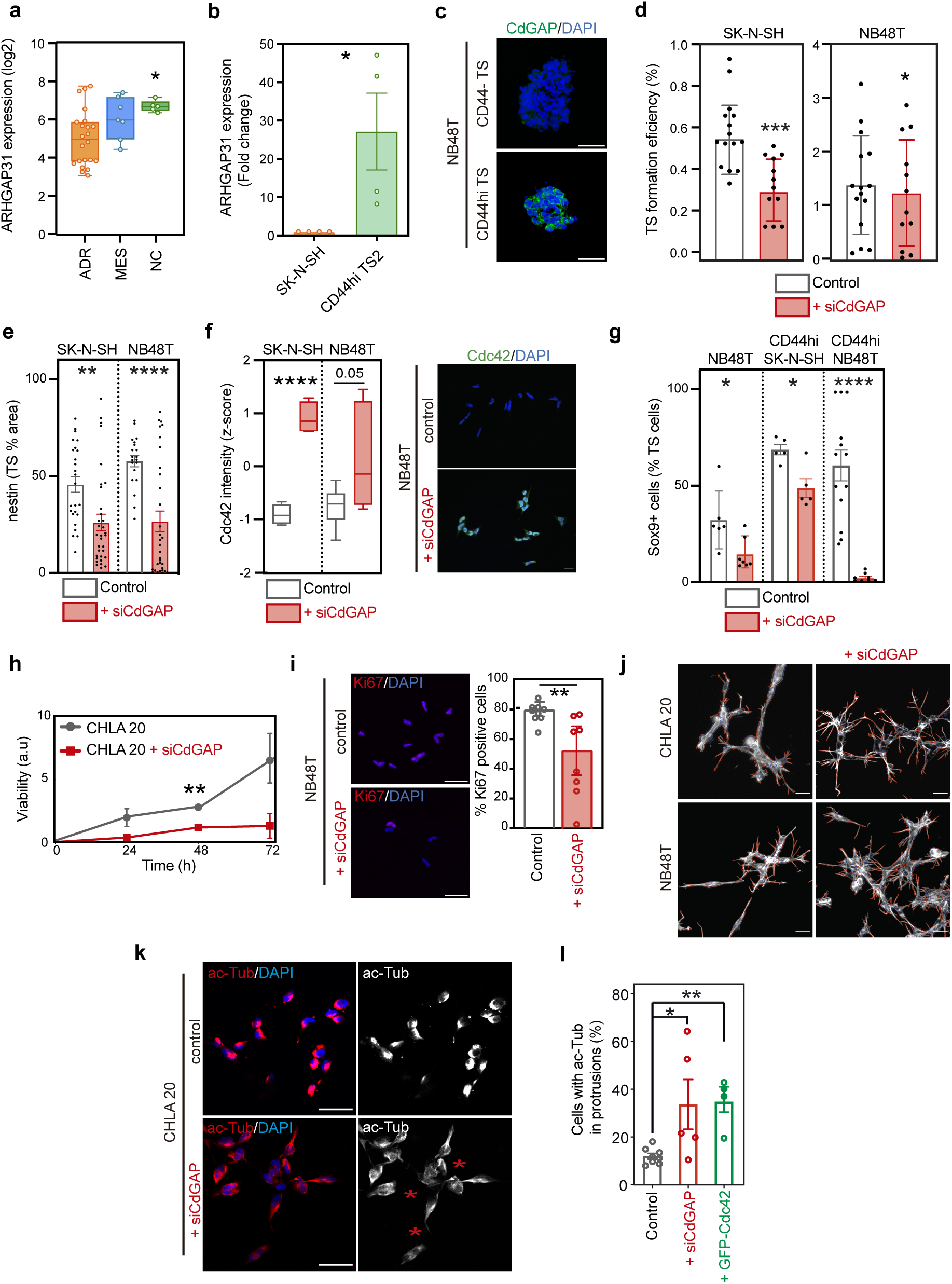
*ARHGAP31*/CdGAP is associated with undifferentiation, and its downregulation provokes Cdc42 overexpression. **a** expression of *ARHGAP31* in adrenergic tumors, mesenchymal tumors or neural crest tissues. Long rank test p value is shown, *=p<0.05. Dataset: GSE908030. **b** Expression of *ARHGAP31* in SK-N-SH CD44hi-derived undifferentiated tumorspheres compared to the bulk cell line. *=p<0.05. **c** Immunofluorescence showing CdGAP expression in NB48T CD44hi-derived undifferentiated tumorspheres compared to CD44neg-derived ones. **d** Tumorsphere formation efficiency of SK-N-SH or NB48T PDX-derived cell lines after CdGAP knock down by siRNA. *=p<0.05; ***=p<0.005. **e** Expression of the undifferentiated marker nestin in SK-N-SH and NB48T PDX-derived cells after CdGAP knock down by siRNA. **=p<0.01; ****=p<0.001. **f** Expression of Cdc42 in SK-N-SH and NB48T PDX-derived cells after CdGAP knock down by siRNA. Representative images from NB48T cells are shown. ****=p<0.001. **g** Expression of Sox9 protein in cells from tumorspheres generated from NB48T or sorted CD44hi SK-N-SH and NB48T cells, after CdGAP knock down by siRNA. *=p<0.05; ****=p<0.001. **h** Cell viability in CHLA 20 cells, with or without downregulation of CdGAP, over 72 hours (**=p<0.01). **i** Quantification of proliferating Ki67 expressing cells in NB48T cells with or without downregulation of CdGAP. Graph shows mean +/−SEM. Scale bar = 20μm. (**=p<0.01, n=2). **j** morphological characterization of CHLA 20 and NB48T cells with or without downregulation of CdGAP. Cell segmentation on the cell contour is shown in red. Scale bar = 20 μm. **k** Immunofluorescence for acetylated tubulin, and DAPI in CHLA 20 cells with or without downregulation of CdGAP. The presence of acetylated tubulin in cell extensions is indicated with red asterisks. Scale bar = 20 μm. **l** Quantification of the cells positive for acetylated tubulin staining in cellular protrusions after the indicated treatments in CHLA20 cells (mean ± SEM, *=p<0.05; **=p<0.001, n=3).

We performed silencing of CdGAP with siRNA to discover that its downregulation is enough to reduce the efficiency of undifferentiated tumorsphere formation and to induce the overexpression of Cdc42 both in SK-N-SH cells and in NB48T PDX-derived cells (Fig. 6d-f). Moreover, the downregulation of CdGAP induces the downregulation of the neural crest marker Sox9 in undifferentiated tumorspheres from diverse origin (Fig. 6g). CdGAP downregulation in NB cells recapitulate the phenotypes observed after Cdc42 overexpression, including a reduction in cell proliferation (Fig. 6h and i), an increase in neuronal morphology and differentiation (Fig. 6j), and an accumulation of acetylated tubulin in neurite-like protrusions (Fig. 6k and l).

These results show that *ARHGAP31*/CdGAP specific upregulation in the undifferentiated progenitor cell population could be a relevant factor in controlling Cdc42 action, halting neuronal differentiation and promoting NB malignancy.

## Discussion

In this work we have performed an extensive study on the expression of the Rho GTPase signaling network in NB tumors. Although various reports have now suggested the importance of Rho GTPase-related genes in neuroblastoma progression and metastasis (11,12,27) a systematic evaluation of the involvement of the different Rho subfamilies and their main regulators and effectors had not been performed to date.

Here we describe that Rho GTPase signaling is widely deregulated in NB patient tumors, correlating with adverse clinical parameters. Although Rho GTPases regulation has classically been described in relation to their GTP/GDP loading, many of the Rho GTPases are also regulated at the transcriptional and posttranslational level, or by mRNA stability (14). Expression is even more relevant in the case of the atypical, constitutively active Rho GTPases (*RHOU*, *RHOV*, *RHOH*, *RHOD*, *RHOF* and the *RHOBTB* and *RND* subfamilies) (28). We have predominantly found low expression of Rho GTPase genes significantly correlating with parameters like high risk, metastasis, relapse, low survival or *MYCN* amplification in different NB patient tumor cohorts. Distinguishably, a high expression of *RHOA*, *RHOD* or *RHOV* is significantly associated with malignancy in tumor datasets. The significant correlations found are for the most part in agreement with changes seen in other cancers. For example, a high *RHOA* expression has been associated with oncogenic progression in a wide range of tumors (reviewed in (29)), whereas RhoB has been attributed a tumor suppressor role and is frequently found downregulated in tumors from diverse origin (30). RhoV is involved in neural crest fate specification and has also been found upregulated in lung adenocarcinoma and breast carcinoma (31). Rnd3 has been found downregulated in hepatocellular carcinoma (32) and Cdc42 expression seem to be context dependent and has been found both downregulated or upregulated in tumors compared to normal tissues, although we observed a clear downregulation associated with malignancy in NB (33). The importance of Rho GTPase proteins in the regulation of morphogenesis, differentiation and proliferation, including neuronal development and neural crest function, makes them relevant targets to be involved in processes mediating NB pathogenesis. For instance, Netrins, Semaphorins, Ephrins, BMP or Notch dependent signaling, all extensively described to be altered in high-risk NB, are also master regulators of Rho GTPase action during embryonic development (34,35). At this point, it remains uncertain whether all the observed changes in the expression of Rho family members play a driver or passenger role in the progression of NB.

Given the complex regulation upstream and downstream of Rho GTPases, and their pleiotropic functions, it is difficult to infer the functional consequences of their changes in expression. A more comprehensive view can be obtained analyzing the correlation with regulators and effectors in patient tumor samples. As mentioned above, Rho GTPase regulators have been already described to be mutated in high risk and relapsed NB, although, like with most mutations in NB, they are not recurrent and therefore their role in malignant transformation is unknown (10,11,36). Furthermore, although the importance of the Rho GTPase family of proteins in cancer progression has been recognized and they have been proposed as relevant therapeutic targets, clinical intervention opportunities are closer when considering the wider Rho GTPase signaling network (37). In an attempt to identify signaling networks with functional relevance in NB, our analysis establishes groups of genes with similar expression patterns along tumors and correlating with clinical variables. Two main groups of genes can be identified, with Group 1 relevantly including *RAC2*, *RHOH*, *ARHGAP4*, *ARHGAP15*, *ARHGAP25* or *DOCK8*, all genes implicated in immune system deficient infiltration in tumors (38). Meanwhile Group2 is enriched in genes associated with neuronal differentiation and includes *CDC42*, *RAC1* and several of their regulators from the *PAK* and *DOCK* families (39,40). Relevantly, the Rho effectors ROCK1 and ROCK2 fall into different correlation groups, indicating a differential involvement in NB, as previously suggested (27). In general, differentiation potential appears like a major feature with importance in NB progression participated by these signaling proteins.

The variability in prognosis among neuroblastoma patients necessitates the deintensification of treatment for low- and intermediate-risk patients and the intensification of therapy for high-risk patients to improve clinical outcomes (41). Consequently, efforts have focused on developing new clinical and molecular risk stratification tools to guide clinical decision-making (42). The strong correlation between gene expression and clinical parameters led us to seek a minimal Rho-related gene signature with prognostic value, further validating the significance of these signaling molecules in neuroblastoma. Not surprisingly, genes included in the survival-predictive signature are located in loci affected by chromosomal aberrations previously identified in high-risk neuroblastoma, for example *ARHGEF10L* (1p36.13), *PLEKGH4B* and *ARHGEF28* (Chr. 5), *PAK1* and *ARHGEF12* (11q) or *ARHGDIA* (17q25.3).

The obtained information can be integrated to identify relevant signaling pathways, participated by the Rho GTPase network, and build connecting networks, including RhoGTPases, activators, inhibitors and, more importantly, effectors with relevance in NB progression. However, works focused on Rho GTPase signaling interactome emphasize the low specificity for some of the regulators, especially for GAPs, with regard to their interaction with different Rho GTPase isoforms (43). Furthermore, their function can be affected by the cellular context or subcellular localization, and compensatory effects between some families of GEFs and GAPs can occur, drawing a complex map. Detailed functional analysis with some of the proteins identified can help unravel their implication in NB biology and the susceptibility of these pathways to therapeutic targeting.

As a proof of concept, we have focused on the GTPase Cdc42, whose downregulation we have found strongly associated with malignancy and poor outcome in NB tumors. The *CDC42* gene is located within the 1p36 chromosome region, frequently deleted in high grade NB, and downregulation of *CDC42* has been associated to *MYCN* amplification in NB tumors (44). However, we have observed a general *CDC42* downregulation associated with malignancy independently of *MYCN*, both in tumors and in cells. We have described that low Cdc42 levels are necessary for NB progression and for the proliferation of adrenergic *MYCN* non amplified NB cell lines in vitro and in vivo. Moreover, we have identified an association of Cdc42 with NB cell adrenergic differentiation via the induction of neuritogenesis and microtubule stability. Cdc42 has been previously implicated in neurite growth (18) it is known to control the microtubule cytoskeleton, and we have found it overexpressed upon differentiation. Interestingly, we have shown that upregulation of Cdc42 levels is enough to drive differentiation of undifferentiated tumorsphere-derived NB cells, and its downregulation promotes a more undifferentiated phenotype, indicating an active role of Cdc42 in inhibiting stemness. Cdc42 activity mediates neural stem progenitors’ function and Cdc42 expression loss in neural progenitor cells gives rise to hyperproliferative lesions in the brain in a mechanism involving Sonic hedgehog (45,46). The downregulation of Cdc42 activity in NB tumors appears to be essential for the malignancy associated with an undifferentiated phenotype. This positions Cdc42 as a putative tumor suppressor in NB linked to differentiation and suggests that Cdc42-activating signaling could serve as a potential differentiating therapy. We have identified a mechanism for Cdc42 downregulation, acting specifically in undifferentiated NB progenitor cells, via the overexpression of the Cdc42 GAP *ARHGAP31*/CdGAP. *ARHGAP31* is a GAP with described specificity for Cdc42 and Rac1 (47). We have found that *ARHGAP31*/CdGAP is specially expressed in neural crest progenitor cells and tumorsphere-derived NB progenitor cells, and that the downregulation of the protein leads to a decrease in stemness with the reduction of Nestin+ and Sox9+ cells. CdGAP has been recently involved in stem cell maintenance in the intestine (48). In NB cells, *ARHGAP31*/CdGAP silencing induces a strong upregulation of Cdc42 levels, recapitulating the low proliferative and differentiating phenotypes observed after Cdc42 overexpression. Although *ARHGAP31*/CdGAP has RhoGAP activity against Cdc42 without necessarily affecting its expression, it has been described to have GAP independent activities in cancer cells (49). We therefore delineate a hypothesis in which Cdc42 activity is essential for NB cell differentiation, and a downregulation of Cdc42 drives a malignant undifferentiated phenotype in NB tumors. CdGAP is specifically upregulated in NB progenitor cells, being a possible culprit for Cdc42 deregulation in this malignant cell population. Differentiating agents have been applied in NB in order to treat minimal residual disease after tumor resection, albeit some patients have manifested resistance to this kind of therapy (50). A deeper understanding of the differentiation-related signaling in NB, such as the CdGAP/Cdc42 axes in undifferentiated cells, could help improve the efficacy of these therapeutic approaches. Although Rho GTPases emerge as important mediators of NB progression, they have proved to be difficult to target (37). The targeting of Rho GTPase regulators such as GAPs and GEFs seems to be a more promising venue and highlight the need to stablish functionally relevant connections among these signaling partners in particular tumors.

## Conclusions

Our study highlights the significant role of Rho GTPase signaling networks in NB progression, with widespread dysregulation of these pathways correlating with poor clinical outcomes. In particular, Cdc42 emerges as a crucial tumor suppressor, whose downregulation by *ARHGAP31*/CdGAP in progenitor cells contributes to the maintenance of an undifferentiated, malignant phenotype in NB. The identification of a Rho-related gene signature with prognostic value suggests that these signaling molecules could serve as important biomarkers for risk stratification in NB patients. Furthermore, the restoration of Cdc42 activity offers a promising therapeutic approach to promote differentiation and inhibit NB progression. These findings emphasize the potential of targeting the broader Rho GTPase regulatory network to develop novel therapeutic strategies for NB.

## Abbreviations

NB: neuroblastoma
GEF: guanine exchange factors
GAP: GTPase activating factors
GDI: guanine dissociation inhibitors
PI: Patient-derived xenograft
ATRA: all-trans retinoic acid
PI: prognosis index

## Declarations

### Ethics approval and consent to participate

All animal experiments were conducted at the facilities of the Instituto de Biomedicina de Sevilla according to procedures approved by the Ethics Committee from the University of Seville and complying with all animal use guidelines. Fresh neuroblastoma tumor samples were managed and obtained from the Andalusian tissue Biobank (Andalusian Public Health System Biobank and ISCIII-Red de Biobancos PT13/0010/0056) after informed consent were obtained from subjects’ guardians, following all established regulations. Clinical information on patient cohorts is available for research use and has been anonymized.

### Availability of data and materials

The datasets supporting the conclusions of this article are included within the article and its additional files.

### Authors’ contributions

MAG substantially contributed to the acquisition, analysis and interpretation of all data in the manuscript. MO contributed to the acquisition of experimental data. LL contributed to the analysis of gene expression data. IR provided technical assistance and acquired data. AA contributed to the cellular profiling and in vivo experiments. JAC contributed to the generation of the gene expression signature and prognostic analysis. MAG, RP and FMV contributed to the conception and design of the work and interpretation of data. MAG, RP and FMV wrote the original manuscript. All authors read and approved the submitted manuscript.

## Acknowledgements

We acknowledge the support of F.J. Morón and R. March from the Genomic’s facility and Rocío Durán from the Histology’s facility at IBiS.

## Additional files legends

**Additional Fig. S1 (related to Fig. 1).** Expression of the Rho GTPase family in relation to different clinical parameters in NB. Box and whiskers representation showing the expression distribution of Rho GTPases in patient tumors according to (**a**) INSS stage of the disease (data from GSE45547 dataset), (**b**) INRG risk classification (data from GSE62564 dataset) or (**c**) MYCN amplification status (data from GSE45547 dataset). *:p<0.05, **:p<0.01, ***p<0.001, ****p<0.0001.

**Additional Fig. S2 (related to Fig. 1).** Expression of the Rho GTPase family in relation to different clinical parameters in NB. Box and whiskers representation showing the expression distribution of Rho GTPases in patient tumors according to (**a**) recurrence in NB patients (data from GSE16476 dataset), (**b**) relapse in NB patients (data from GSE3446 dataset) or (**c**) histology (data from TARGET dataset). Only the GTPases with significant correlations are shown. *:p<0.05, **:p<0.01, ***p<0.001, ****p<0.0001.

**Additional Fig. S3 (related to Fig. 1).** Patient tumors stratification according to Rho GTPase expression. K-means clustering of patients according to Rho GTPase expression in the GSE16476 (**a**) and GSE45547 (**b**) datasets. Age at diagnosis, survival, INSS stage and MYCN amplification status are shown.

**Additional Fig. S4 (related to Fig. 1).** Kaplan-Meier overall survival analysis based on Rho GTPase expression. Overall survival did not take into account relapsed events. Data from cohort GSE62564. n= number of samples. Bonferroni adjusted p values are shown.

**Additional Fig. S5 (related to Fig. 2).** Rho signature prediction validation**. a** and **b** Area under the curve (AUC) analysis of the predictive capacity of the Rho gene signature on the overall survival of NB patients during the first three years from diagnosis (t = time in days, FP = false positives or specificity, TP = positives or sensitivity). (a) presents data from dataset TARGET and (b) from GSE62564. **c** Kaplan-Meier analysis using the prognostic index (PI) of the Rho gene signature. Samples with higher PI values have lower survival than the group of samples with lower PI (Strata: classification of patients in survival data according to PI value, PI High = high PI values, PI Low = low PI values) **d** Hazard ratio (HR) obtained from a Univariate Cox Analysis for the PI value and other clinical parameters in relation to the overall survival of the patients (p value = Likelihood ratio test, Wald test and logRank test). Data from cohort GSE62564. **e** STRING analysis of proteins with the greatest possibility of interaction with the genes in the signature. The network representation with proteins as nodes together with the protein-protein associations are shown. Protein-protein interaction enrichment p-value < 1×10^−6^.

**Additional Fig. S6 (related to Fig. 3). a** quantification of Cdc42 signal intensity from images described in Fig. 3A. (log a.u= log of arbitrary units, **=p<0.001). **b** western blot showing the results of GTPase pull down assay demonstrating the activity of the expressed GFP-Cdc42 in CHLA 20 cells. PD: pull down. C: control. **c** Graphs showing tumor onset and growth of CHLA 255 in orthotopic xenografts with or without the expression of GFP-Cdc42. LogRank statistical analysist significance is shown. Ns: not significant. Mean +/− SD is plotted in cell growth graph. **d** Immunohistochemistry from tumors described in C showing the expression of the indicated markers. Quantification shows the % of positive cells in each condition (mean +/− SEM, *=p<0.05). Scale bar = 50 μm.

**Additional Fig. S7 (related to Fig. 6)**. The Cdc42 regulator *ARHGAP31*/CdGAP is highly expressed in undifferentiated neural crest-like NB cells. **a** Heatmap representing the expression of genes from the Rho GTPase signalling network in tumor and cell lines with adrenergic, mesenchymal or neural crest character. ADR: adrenergic; MES: mesenchymal; NC: neural crest cells or cell lines. Data from GSE908030 cohort. The lists of genes with a higher expression in MES and NC or in ADR are shown. **b** Western blot showing the downregulation of *ARHGAP31*/CdGAP after transfection with an specific siRNA in the CHLA20 cell line. **c** *ARHGAP31*/CdGAP relative expression levels in siRNA treated CHLA20 and NB48T cell lines compared to controls. **d** *ARHGAP31*/CdGAP relative expression levels in siRNA treated SK-N-SH cells compared to controls. f.c.: fold change.

**Additional Fig. S8**. Full uncropped blots.

**Additional Table S1:** Patient cohort databases used in the study.

**Additional Table S2:** Rho-related selected genes used in the study.

**Additional Table S3:** DNA primers used in the study.

**Additional Table S4.** Results from the expression analysis on Rho-related network genes. Complete results from the Pearson correlation analysis (GSE45547 dataset) and the gene cluster resulting are shown. The complete list of genes comprising the different gene modules resulting from the WCGNA analysis and the gene groups from the JACCARD analysis are shown on different tabs. GO and Reactome enrichment analysis are also shown for the groups 1 and 2.

