## Supplementary figures and images for "Rho GTPases signaling mediates aggressiveness and differentiation in neuroblastoma tumors"

### Supplementary Figures S1 to S8

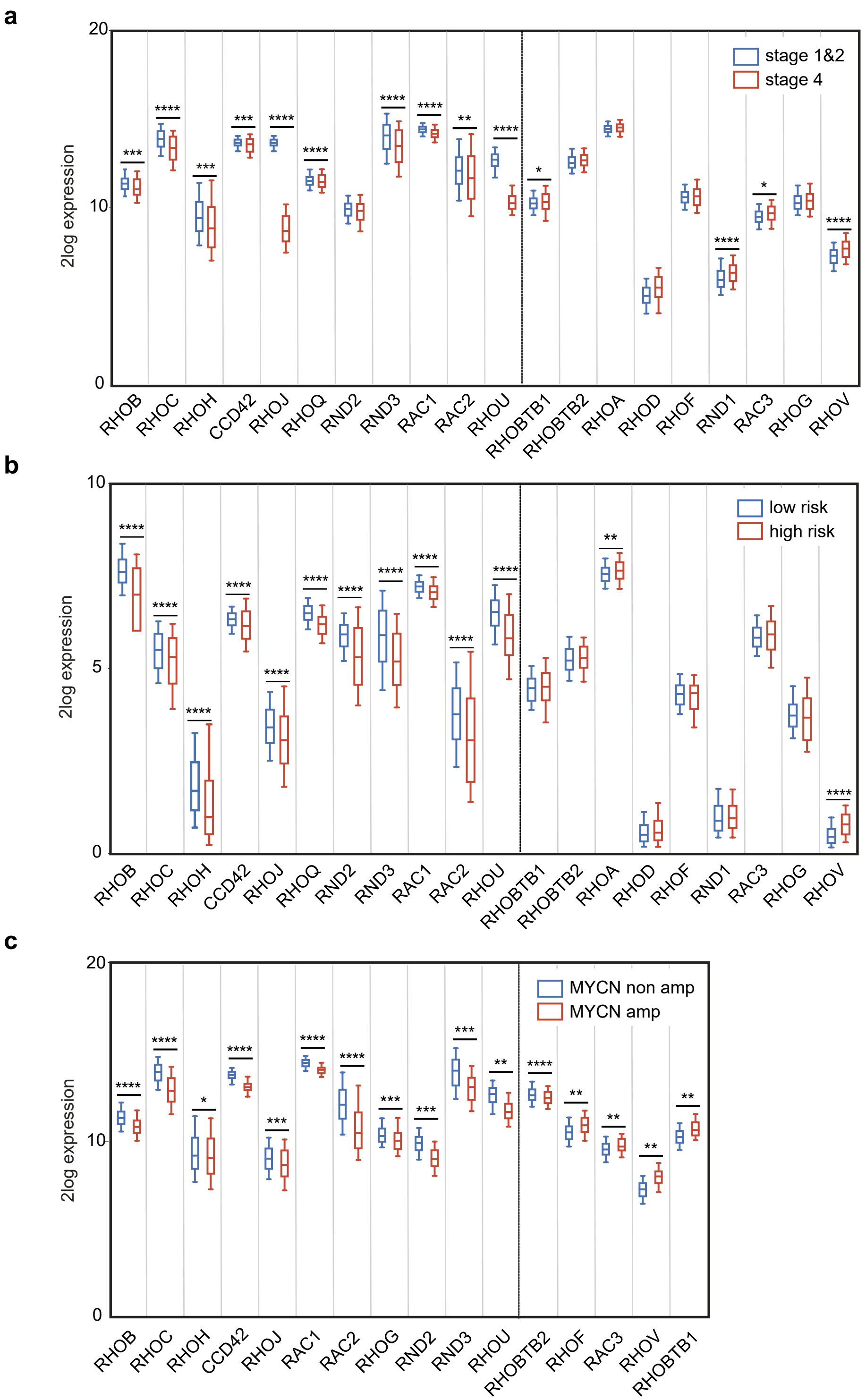

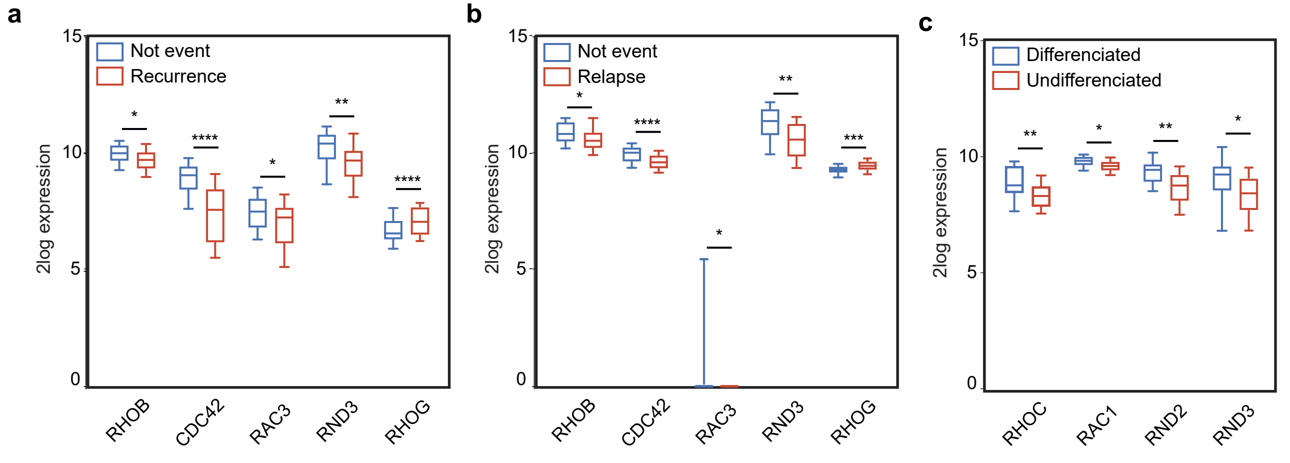

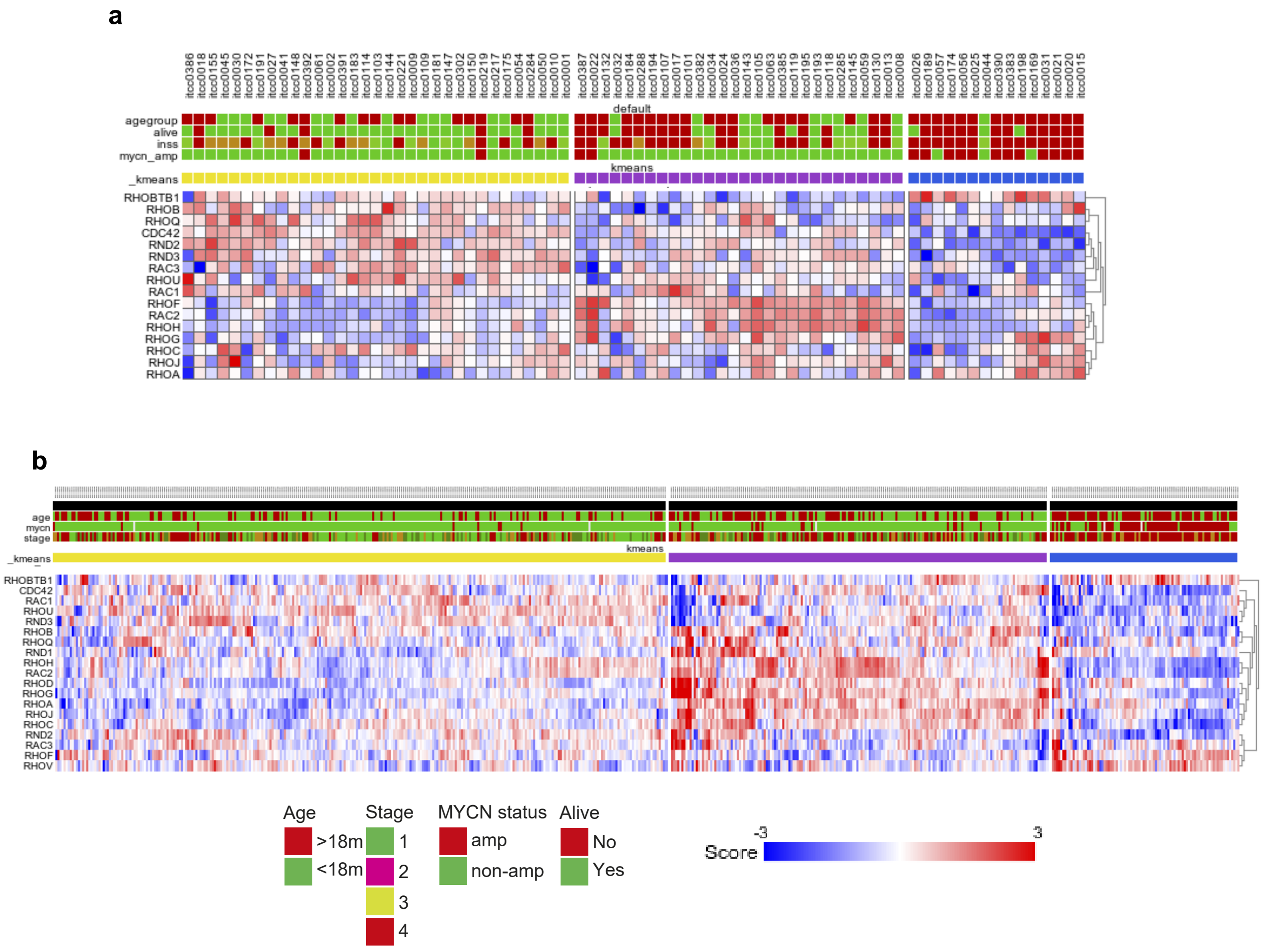

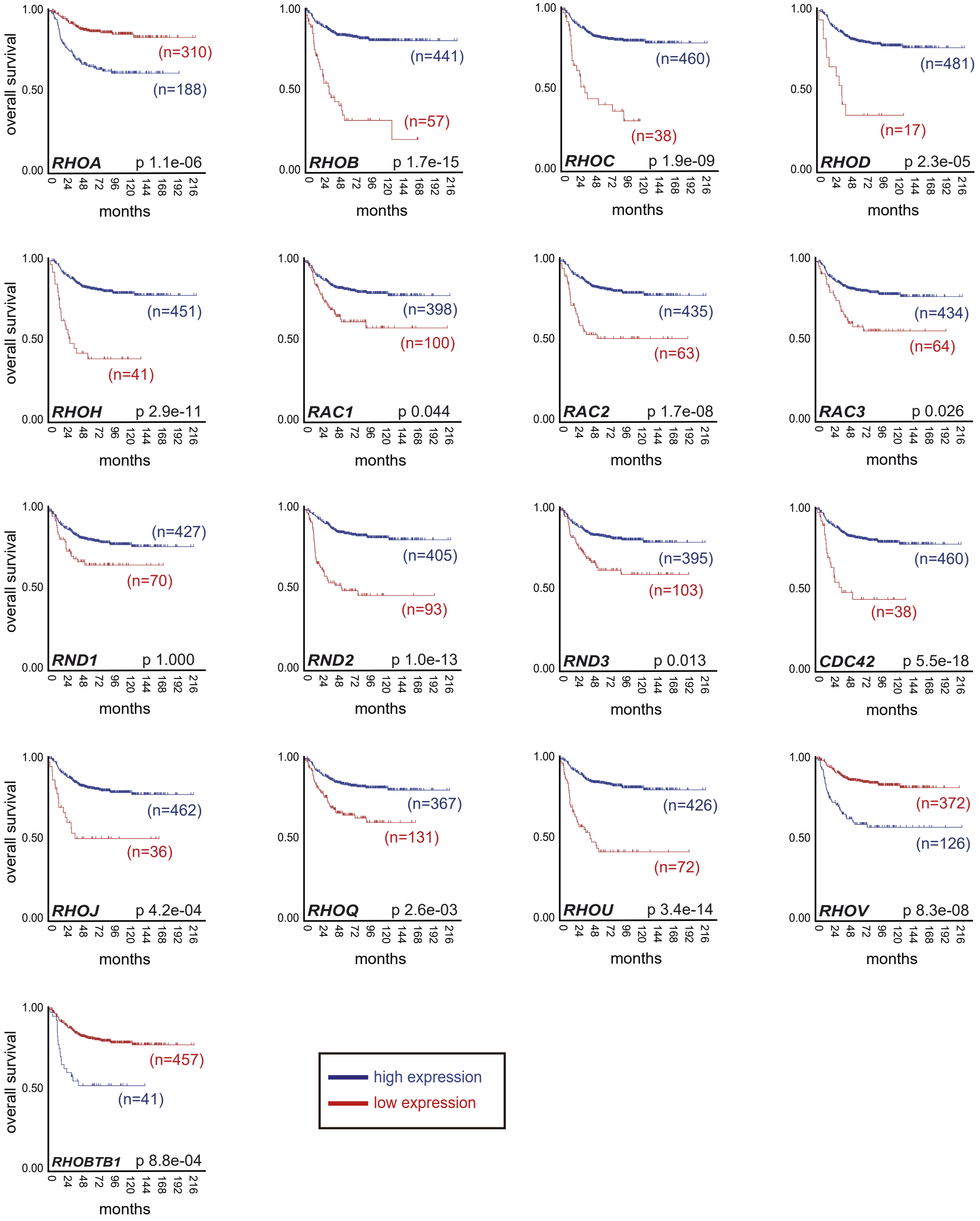

**Figure S5**

a

b

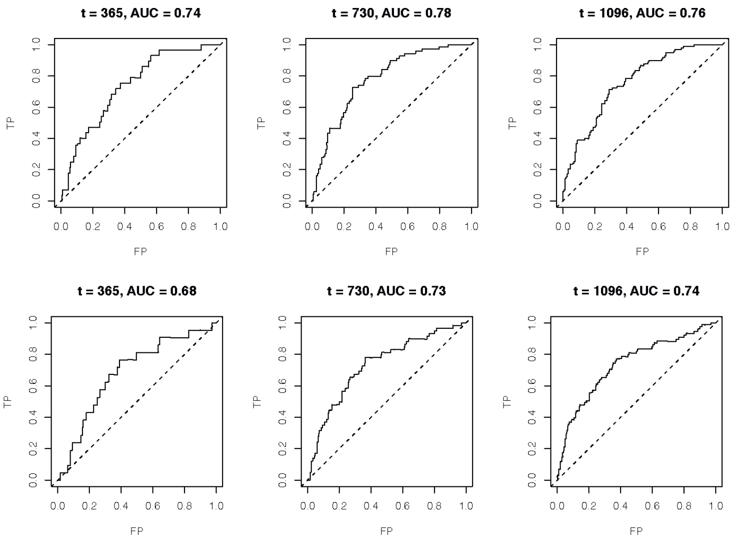


c

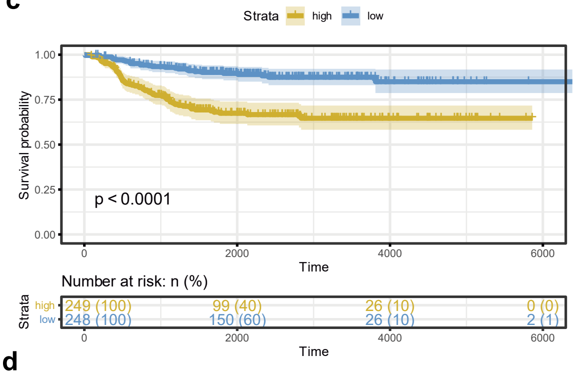


d

| | beta | HR | HR_LCI | HR_HCI | wald.test | p.value |
|---|---|---|---|---|---|---|
| Age | 0.00034 | 1.0 | 1.0 | 1.0 | 62 | <0,0001 |
| MYCN | 210.000 | 7.8 | 5.3 | 12.0 | 100 | <0,0001 |
| PI | 0.23000 | 1.3 | 1.2 | 1.4 | 36 | <0,0001 |

e

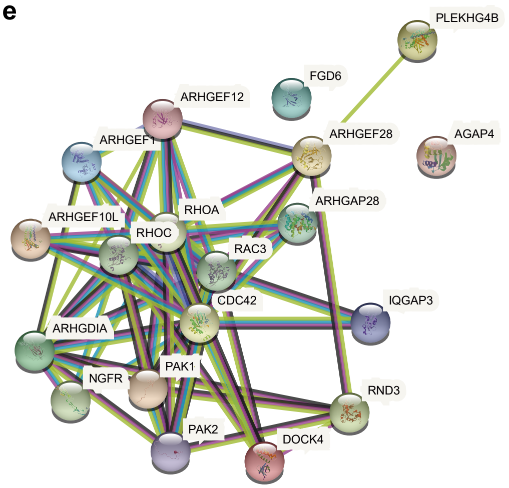

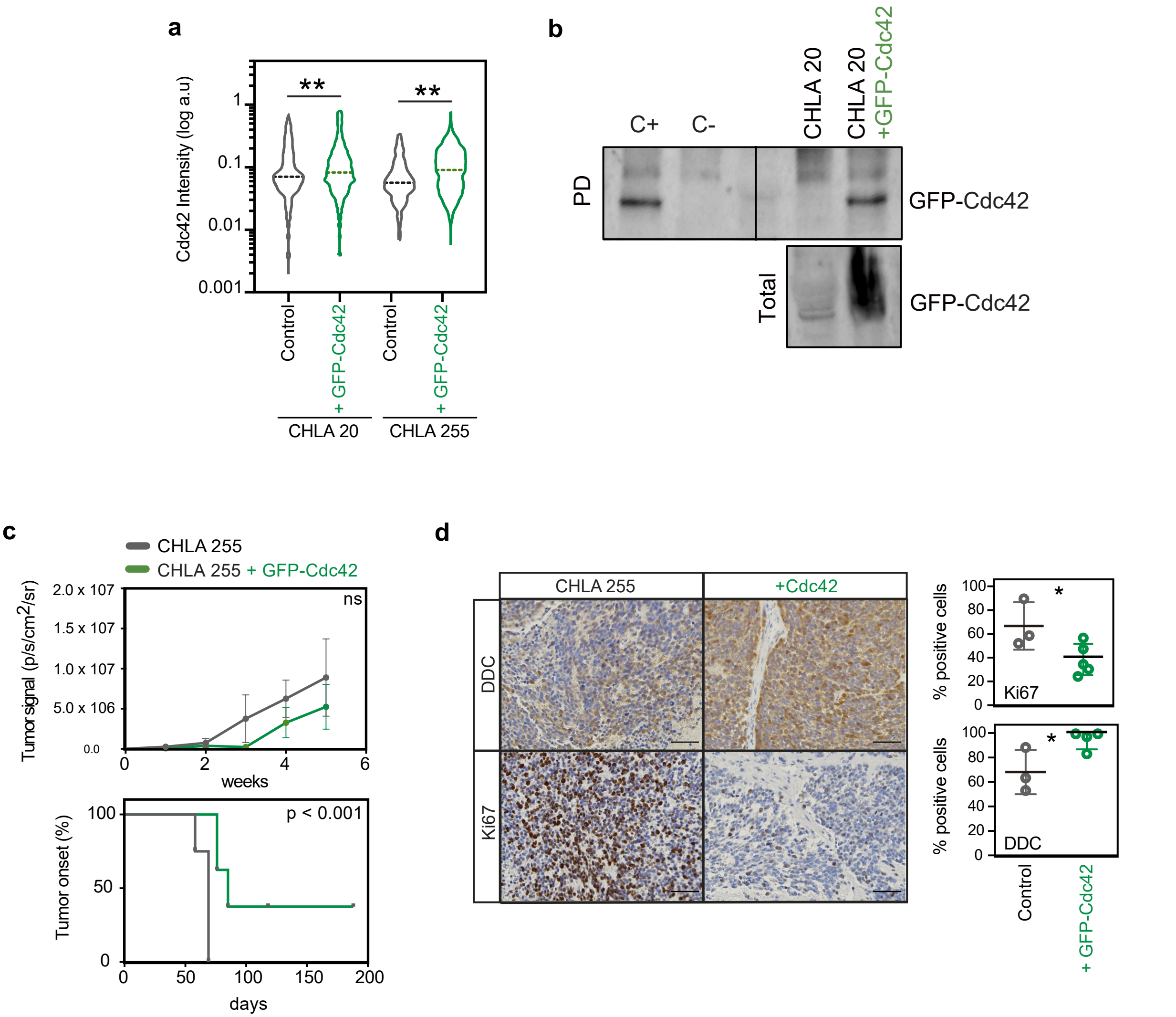

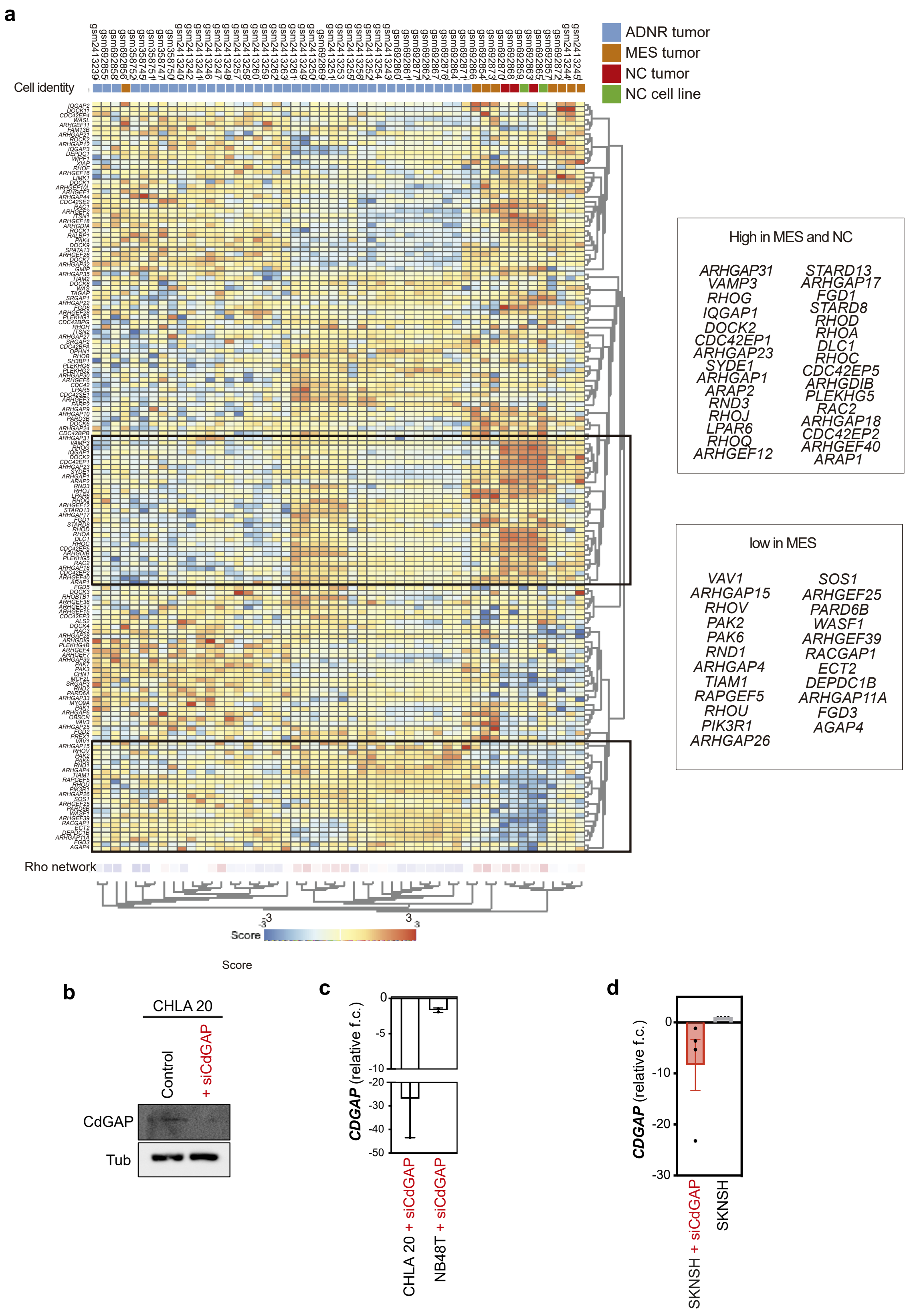

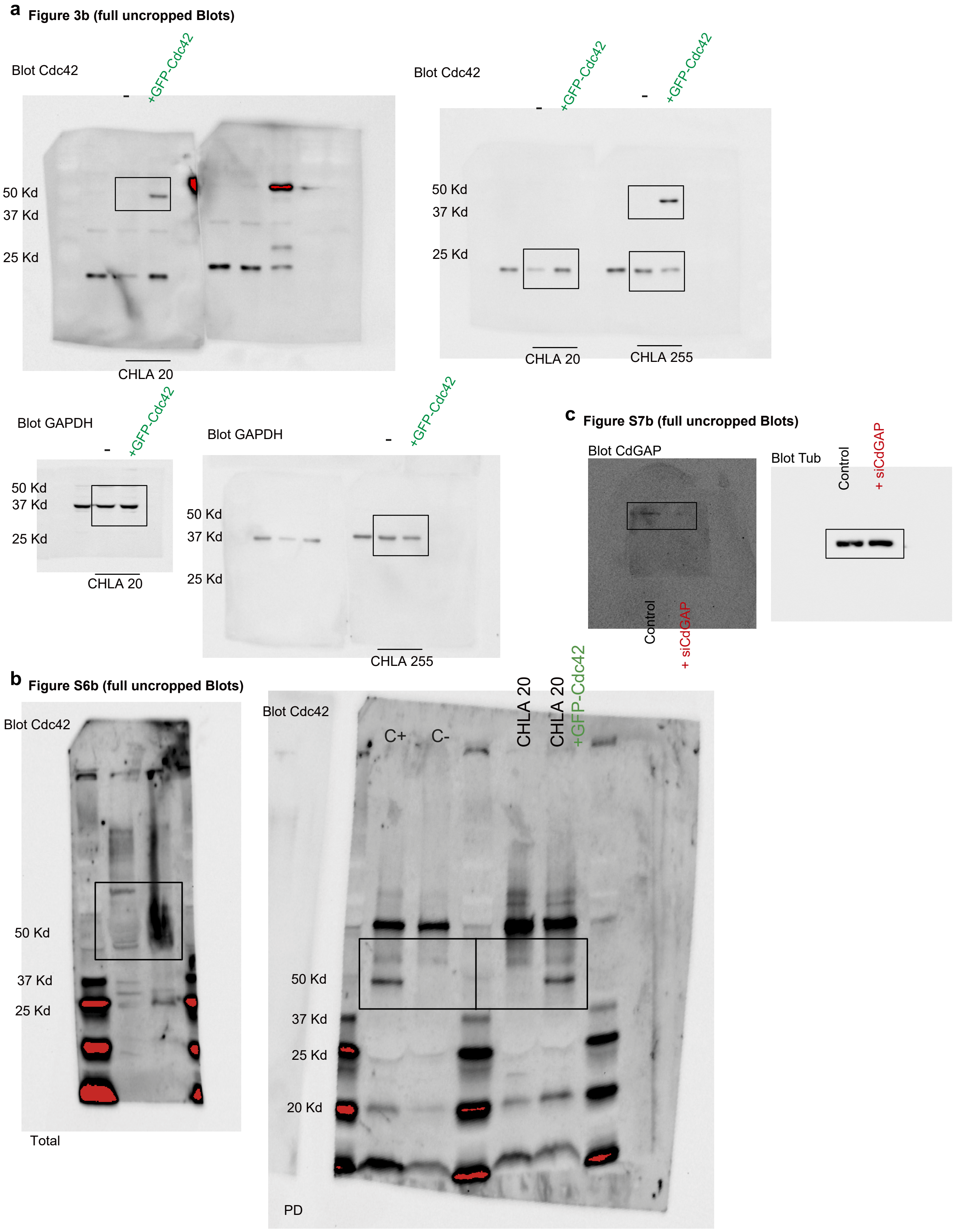
